# Paternal ancestral gestational exposure to neonicotinoid thiacloprid induces transgenerational alteration in DNA methylation germline germline-specific genes in female ovaries

**DOI:** 10.1101/2025.07.29.665494

**Authors:** Ouzna Dali, Chaima Diba Lahmidi, Christine Kervarrec, Pierre-Yves Kernanec, Fatima Smagulova

**Author notes:** Corresponding author Fatima Smagulova, Irset-Inserm UMR 1085, 9 avenue du Prof. Léon Bernard, 35000 Rennes, France. Ouzna Dali, Chaima Diba Lahmidi, Christine kervarrec, Pierre-Yves Kernanec, Fatima Smagulova.

## Abstract

**Background:** Neonicotinoids, a widely used class of insecticides, have raised concerns due to their potential role in the decline of bee populations. Accumulating pieces of evidence suggest that these chemicals may also pose risks to human and animal health.

**Objectives:** This study investigates the transgenerational effects of thiacloprid (Thia), a neonicotinoid, on the female reproductive system.

**Methods:** Pregnant outbred Swiss female mice were exposed to 0 and 6 mg/kg/day of thiacloprid from embryonic days E6.5 to E15.5. Adult female F1 and F3 progeny ovaries were examined for morphological abnormalities by using hematoxylin and eosin (H&E) staining of paraffin sections. Additionally, estradiol levels were assessed by ELISA using blood serum. A marker of mitosis, phosphorylated histone H3 at serine 10 (PhosphoHisH3ser10) levels, was analyzed by immunofluorescence, DNA methylation at some regions was analyzed by MeDIP, and RT-qPCR assessed gene expression.

**Results:** The gestational exposure to thiacloprid led to an increased formation of multi-oocyte follicles in directly exposed F1 generation ovaries but not in F3. We observed a reduction in serum estradiol levels in both F1 and F3 females. In direct F1 exposed progeny, we determined an increase in (PhosphoHisH3ser10) in granulosa cells. Additionally, alterations in DNA methylation were observed in germline reprogramming-responsive genes in embryonic ovaries, as well as the DNA methylation was altered at these genes in both the F1 and F3 ovaries. The modified DNA methylation was associated with changes in gene expression in adult ovaries of the F1 and F3 generations in genes encoding transcription and hormonal signaling factors.

**Conclusion:** Our findings demonstrate that ancestral gestational exposure to thiacloprid induces transgenerational effects on female reproductive health, potentially through epigenetic modifications of paternal germ cells.

## Introduction

The widespread use of neonicotinoid pesticides has raised significant concerns about their potential impact on human and animal health. Among these, thiacloprid is a commonly used neonicotinoid valued for its effectiveness against diverse agricultural pests [1]. However, its environmental persistence and emerging evidence of toxicity raise serious concerns about its safety for mammalian species, particularly during critical, vulnerable developmental periods [2]. This study is a continuation of an integrative study of our laboratory, where we investigated the impact of thiacloprid on mammalian organs. Specifically, previously we showed that ancestral exposure to thiacloprid causes alterations in the thyroid gland [25], male reproductive organs, testis [26], and prostate (Dali et al, submitted). This study aims to reveal the effects on the female ovarian system. Prenatal exposure to environmental chemicals is particularly vulnerable due to epigenetic reprogramming events and the formation of many organs. Among them, the ovary, a key organ of the female reproductive system, is highly sensitive to endocrine-disrupting compounds (EDCs). EDCs can interfere with ovarian function by mimicking or blocking hormones, disrupting hormonal signaling, and affecting hormone synthesis. This interference can lead to various reproductive health issues, including altered ovarian development and function [27]. The epigenetic reprogramming and initiation of prophase I of meiosis during fetal development make ovaries very fragile, with potential long-term consequences for fertility and endocrine function [3]. Research has shown that endocrine-disrupting chemicals can impair follicular development, alter steroidogenesis, and induce oxidative stress in ovarian tissue [4].

It has been shown that oral administration of thiacloprid at a high dose of 50 mg/kg/bw during pregnancy in female rats led to a significant increase in levels of 17β-Estradiol, prostaglandins F2α and E2, and serum oxytocin hormone in different stages of pregnancy, whereas, progesterone and endothelin-1 serum levels have decreased. The authors noted a substantial rise in lipid peroxidation levels [43].

Despite these findings, the impact of prenatal thiacloprid exposure at relatively low dose on ovarian development and epigenetic regulation of reproductive genes remains poorly understood.

This study aims to address this gap by investigating the effects of prenatal thiacloprid exposure on ovarian morphology, follicular development, and estradiol levels in mice. Additionally, we explore potential epigenetic changes, such as DNA methylation, which may contribute to ovarian dysfunction. By examining these outcomes, this research seeks to provide valuable insights into the broader implications of neonicotinoid exposure for reproductive health, with a focus on translational relevance to human fertility risks [5].

## Results

### Gestational exposure to thiacloprid leads to a decrease in body weights in F3 females and to alterations in the ovary-to-body weight ratio in F1 and F3

In this study, we exposed pregnant females to thiacloprid or vehicle from embryonic day E6.5 to E15.5, and female progeny were sacrificed at the age of 2 months; we called them F1 females and these are directly exposed animals as reproductive organs in females are formed during embryonic period. F3 females were paternally exposed animals, as it was part of our previous studies, and many F3 females were generated. In addition, the impact of ancestral paternal exposure on the F3 female reproductive system is not well studied. To get transgenerationally exposed females, the males of F1 exposed progeny were crossed with unexposed females, and F2 males were generated. Next, F2 males were crossed again with unexposed females to obtain the F3 generation. The females of F3 generations were sacrificed at the age of 2 months, and the ovaries and blood were preserved.

To assess the effect of thia exposure on body weight, we weighed F1 and F3 female mice (Fig. 1a). No significant difference in body weight was observed in F1, but a significant decrease (p = 0.01) was noted in F3 females exposed to thia. Additionally, the ovary-to-body weight ratio in thia-exposed mice compared to controls increased in F1 (p = 0.017), while it decreased in F3 (p = 0.05) (Fig. 1b)

**Fig 1.**
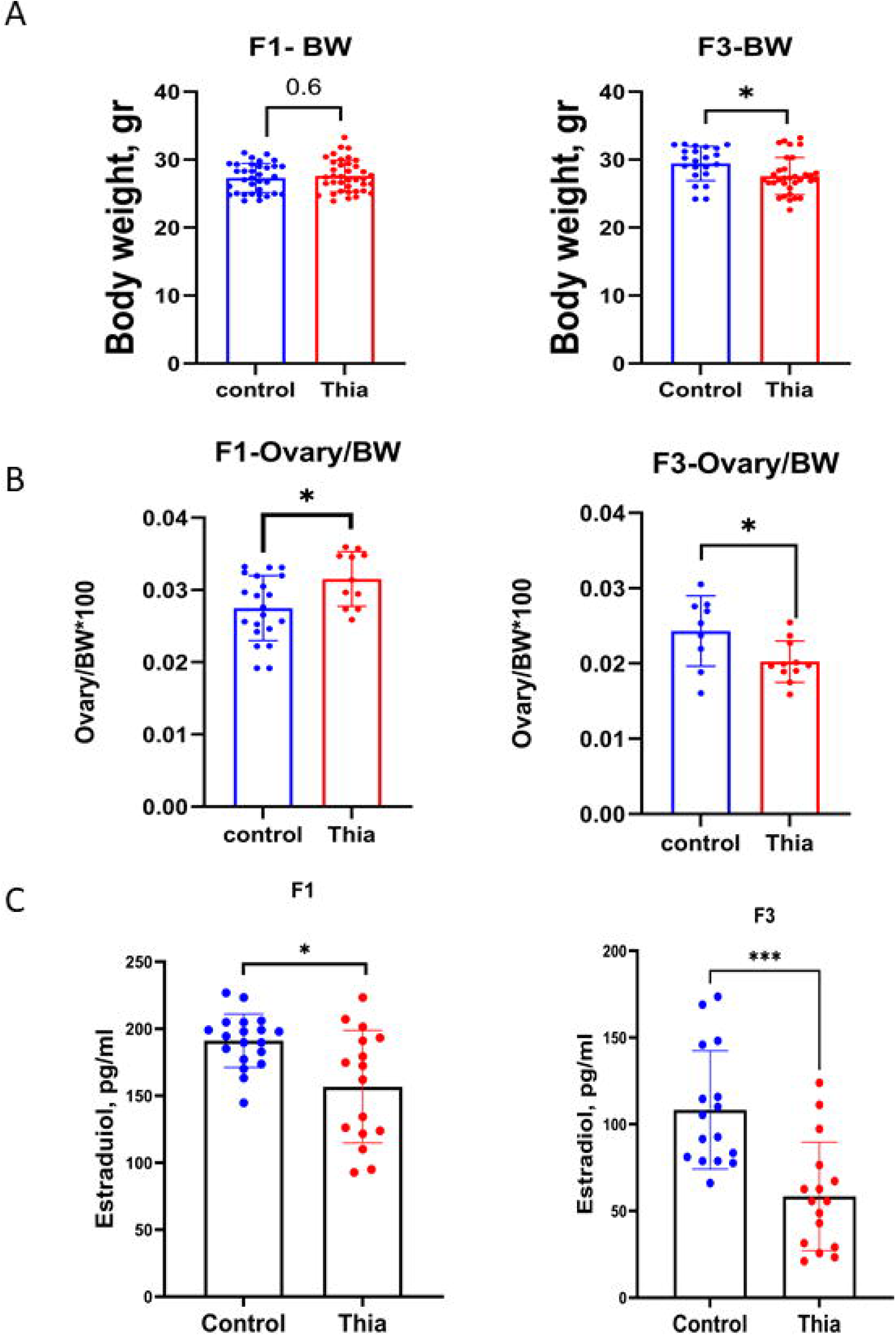
Effects of Thia Exposure on Body and Ovary Weight Across Generations. (A) Body weight changes in F1 and F3 generations. (B) Ovary weight in F1 and F3 generations. (C) Serum estradiol levels in F1 and F3 generations.

Our study suggests that direct exposure in F1 and paternal exposure in F3 affected the female reproductive system.

### Gestational exposure to thiacloprid impacts estradiol levels in the blood serum

Since estradiol is primarily produced by the ovaries, we investigated the impact of gestational exposure on the blood serum level of estradiol. We performed analysis using blood serum and ELISA kit as described in the Methods section. The 100 ul of blood serum or standard dilutions was incubated with the antibody, and an ELISA test was performed according to the instructions provided by the manufacturer. The analysis showed that the thia groups of animals had a reduction of estradiol serum levels in both F1 and F3 generation females. Specifically, in F1, estradiol levels were significantly decreased in the treated group by 0.8-times (p = 0.015), and by 0.5-fold in F3 (p = 0.0001) (Fig. 1c).

Our data suggest that, similar to males, females also have alterations in sex hormones due to the direct or indirect endocrine-disrupting properties of thiacloprid.

### Morphology studies showed a tendency to increase in atretic follicles and an increase in multi-oocyte follicles in F1

To assess the impact of prenatal exposure to thiacloprid on the follicular population, we analyzed histological sections of ovaries from F1 generation mice. Follicle types were counted as outlined in the Methods section. Statistical analysis revealed a tendency towards an increase in atretic follicles by 1.4-times and a significant increase in multi-oocyte follicles (MOF) in F1 (p = 0.04) (Fig. 2d-2e). Further comparisons between the F1 and F3 generations were made to evaluate the potential long-term effects of prenatal exposure by counting the number of MOF in F3 (Fig. 2e).

**Fig 2.**
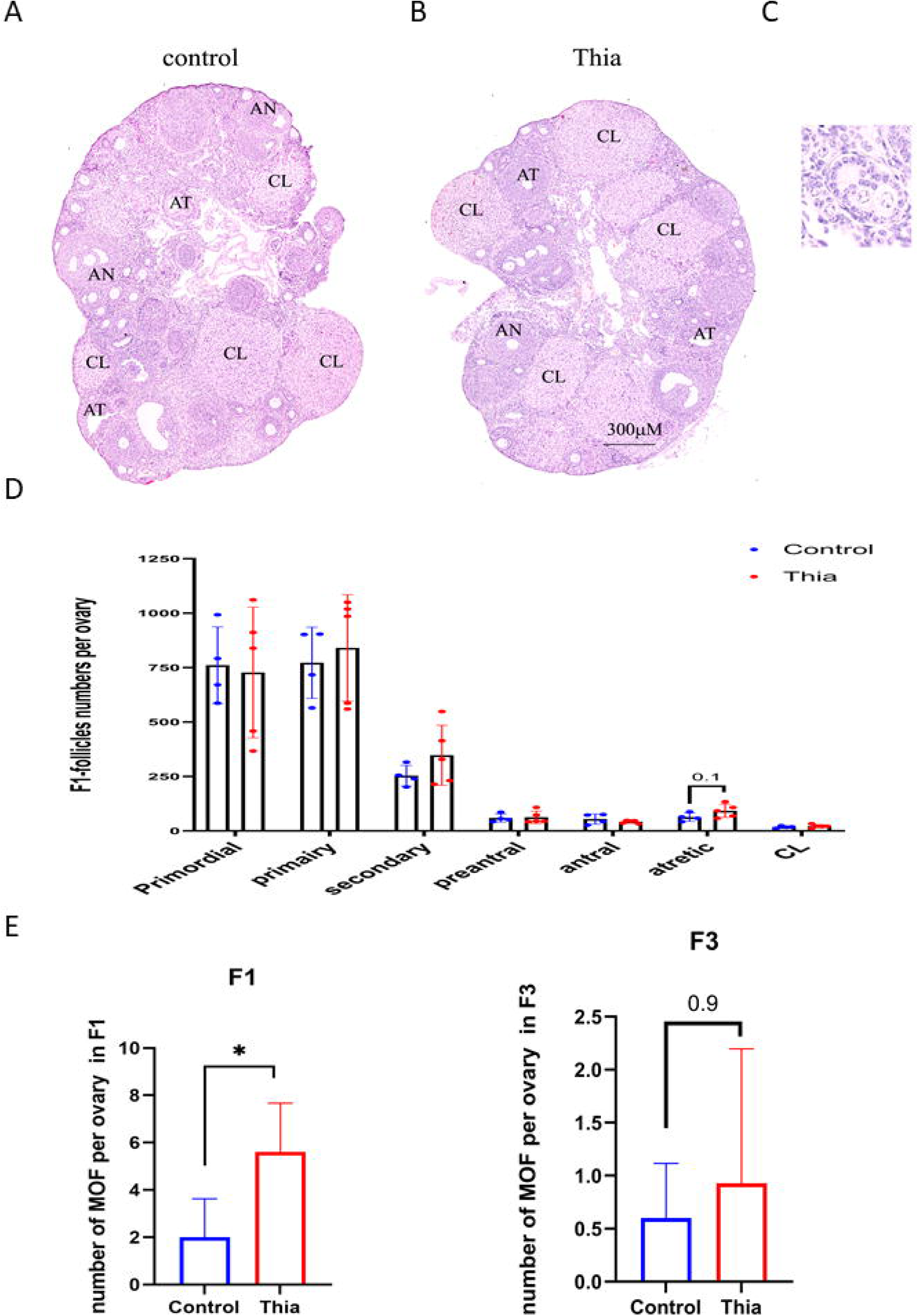
(A, B) Representative H&E-stained ovarian sections from control and Thia exposed groups, respectively. In control ovaries, most late-stage follicles appear healthy, whereas in treated ovaries, numerous atretic follicles are observed, (C) Multi-oocyte follicle (MOF) (NanoZoomer, 5× magnification). (D) Quantitative analysis of follicle types in F1, manually counted using NDP.view2 software by two independent researchers and categorized according to the classification in the Methods section (n = 4 control, n = 5 treated). (E) Quantitative analysis of MOF in F1 and F3. Statistical significance was determined using the nonparametric Mann-Whitney test.

**Fig 3.**
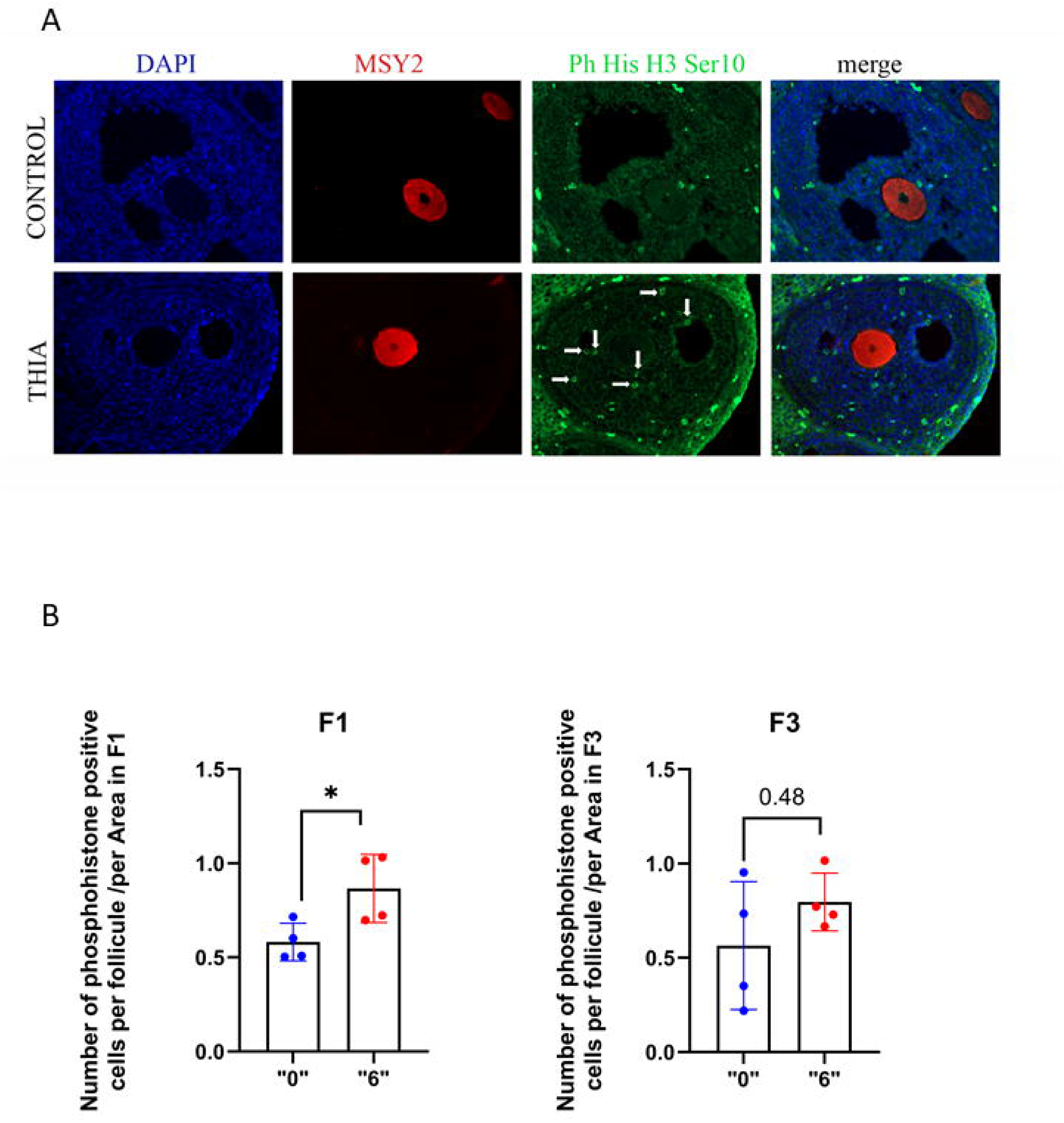
Immunostaining of Phospho-Histone Ser10 and MSY2 in Control and Thia-Exposed Ovaries. (A) Representative images of control and Thia-exposed ovaries. Arrows indicate positive cells. (B) Quantification of phospho-histone-positive cells in F1 and F3 generations. Statistical analysis shows p-values of 0.04 and 0.27, respectively, nonparametric Mann-Whitney test.

Thus, our analysis reveals that morphologically, ovaries were not dramatically affected in F1.

### The mitosis marker PhosphoHisH3ser10 level has increased in directly exposed F1 ovaries

The decrease in estradiol (E2) levels, a form of estrogen, and the increase in multi-oocyte follicles (MOF) could negatively impact ovarian function, particularly by impairing the mitosis of granulosa cells and thereby impacting the oocyte growth and maturation. To investigate this, we analyzed phosphorylated histone H3 levels at serine 10 PhosphoHisH3ser10 using immunofluorescence technique on paraffine sections of the ovaries. We coimmunostained the ovaries against phosphorylated histone H3 at serine 10 and the marker of oocyte MSY2. MSY2 appeared as a strong cytoplasmic marker in oocytes, while PhosphoHisH3ser10 is appeared only at granulosa cells as bright staining all over the nucleus. We analyzed the number of positive cells per antral oocyte. Our results revealed a significant 1.48 times increase in phosphorylated histone H3 positivity in adult F1 ovaries (p = 0.04) following paternal gestational exposure to thia. However, in F3 ovaries, the p-value of 0.27 (t-test) indicates that this trend is not statistically significant. These findings suggest that the thia exposure may perturb the mitosis and thereby alter the function of granulosa cells, which could contribute to the alteration of the production of estradiol and ultimately to correct follicle maturation.

### DNA methylation was increased at germ cell-reprograming-responsive genes

Considering the observed disruptions in follicle development, we hypothesized that prenatal exposure to thiacloprid might induce epigenetic changes during the somatic-to-germline transition. This process involves the global loss of somatic cell DNA methylation marks and the establishment of a new pattern of germline-specific epigenetic marks. To investigate whether the gestational exposure to thia affects the reprograming, we analyzed the methylation status of germcell-reprograming-responsive (GRR) genes, which are known to be essential for the formation of population of germ cells need [44]. To this end, we dissected embryonic E15.5 ovaries, extracted DNA, and performed MeDIP as described in the Methods section. Given the critical role of DNA demethylation at master regulator GRR genes such as *Ddx4, Hormad1, Tdrd1 and Brdt,* in maintaining the germ cell lineage, we examined DNA methylation at their promoters as well as we looked at the DNA methylation at the promoter of retroelement Line 1, as it is generally highly methylated. Our analysis revealed the increased level of DNA methylation by 1.74-, 1.89-, 1.83-, and 1.73-times in *Brdt, Ddx4, Hormad1, and Stk31, respectively,* in the F1 generation (Fig. 4a). These findings suggest that prenatal thiacloprid exposure may disrupt the epigenetic reprogramming of germline cells, potentially altering the gene expression of corresponding genes, that could contribute to ovarian development and function. To further follow the fate of DNA methylation in adult ovaries, we performed the analysis in F1 adult ovaries.

**Fig 4.**
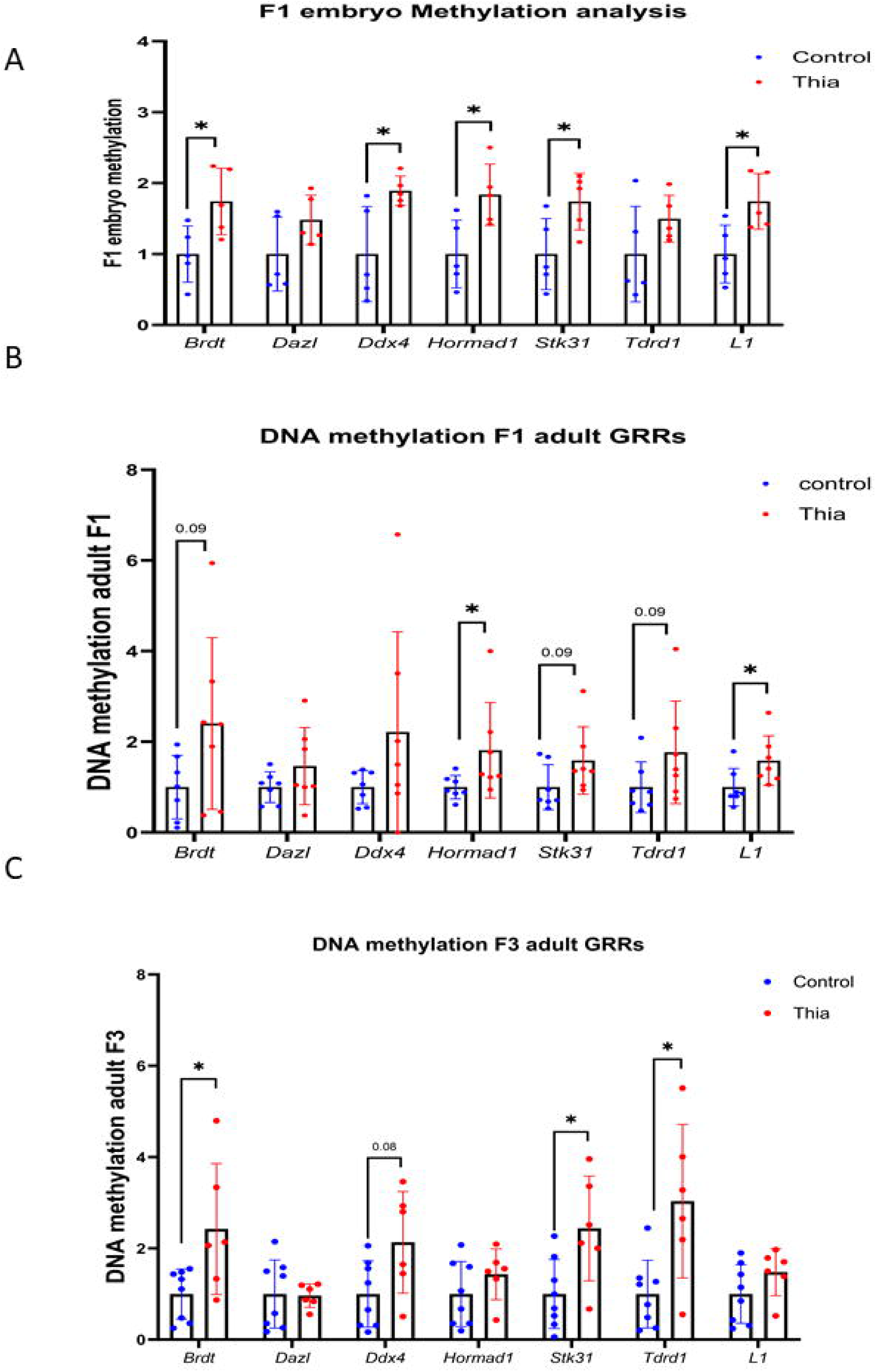
Gestational Thia Exposure Increases DNA Methylation in GRR Genes Across Generations. (A) DNA methylation analysis of GRR genes in embryonic ovaries. (B) DNA methylation analysis in F1 adult ovaries. (C) DNA methylation analysis in F3 adult ovaries.

In F1 adult ovaries, we observed an increased DNA methylation at *Hormad1* by 1.81-fold in F1 and a tendency to increase in *Brdt, Stk31, and Tdrd1* by 2.4-, 1.58-,1.76-times.

In F3, paternally exposed adult ovaries, we observed a similar pattern, the increase in DNA methylation at *Brdt, Stk31 and Tdrd1* by 2.42-, 2.43- and 3.03- times and a tendency to increase in methylation at *Stk31* and *Tdrd1* was observed.

Our data suggests that prenatal exposure alters DNA methylation in directly exposed female embryonic ovaries and these changes were persistent in adult F1 ovaries. Since F3 generation females were from paternal exposure, these data suggest that paternal DNA methylation could impact future embryonic DNA methylation in female ovaries via altered methylation of sperm, which was previously determined in our study need [45].

### The effects of thiacloprid on genes encoding for gonadal and reproductive regulation

To assess the effect of gestational exposure on gene expression in the adult ovaries we extracted RNA from ovaries and performed RT-qPCR gene expression analysis to evaluate the expression of several candidate genes involved in the estrogen signaling pathway and the transcriptional regulation of ovarian development, including *Inha*, *Inhba*, *Inhbb*, *Foxl2*, *Foxo3*, and *Foxo4*.

Our results revealed significant alterations in gene expression due to prenatal exposure to thia. In the F1 generation, the expression of *Inhba* (p = 0.004) and *Fst* (p = 0.006) was elevated by 2.54-and 0.44-times respectively, In the F3 generation, we observed a more widespread upregulation, with significant increases in the expression of *Inha* (p = 0.02), *Inhbb* (p = 0.05), *Foxl2* (p = 0.02), and *Foxo3* (p = 0.06) 1.59-, 1.83-, 1.45-and 1.52-times (Fig. 5).

**Fig 5.**
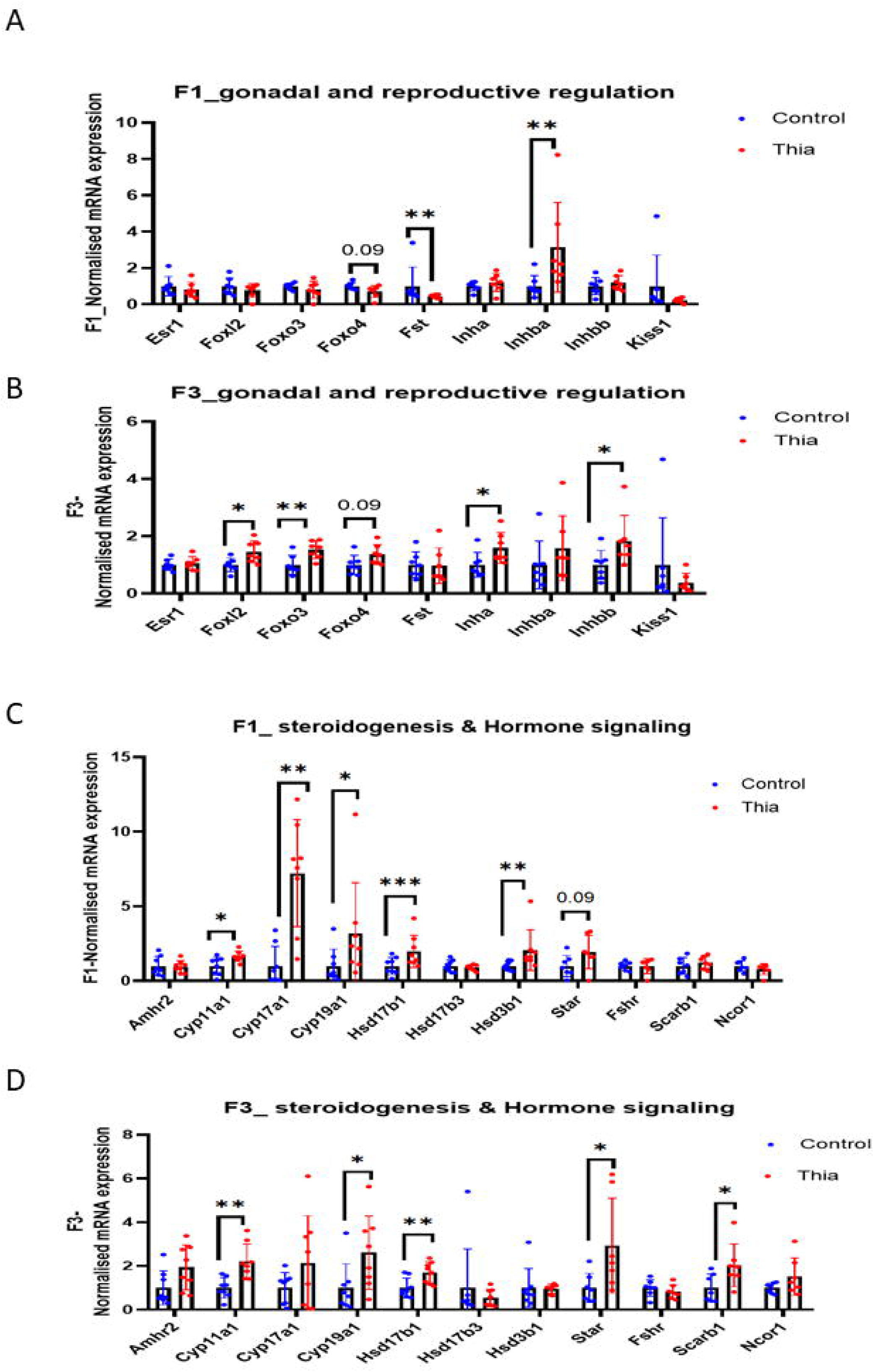
Gestational Thia Exposure Alters Gene Expression in F1 and F3 Ovaries. (A) Expression of gonadal and reproductive regulation genes in F1 ovaries. (B) Expression of gonadal and reproductive regulation genes in F3 ovaries. (C) Expression of steroidogenesis and hormonal signaling in F1 ovaries. (D) Expression of steroidogenesis and hormonal signaling in F1 ovaries. Gene expression levels were normalized to the housekeeping gene *Rpl37a*. Statistical significance: *p* < 0.05 (*), *p* < 0.01 (), *p* < 0.001 (*), *t*-test.

### The effects of thiacloprid on genes encoding for steroidogenesis and hormonal signaling

To assess the impact of gestational thia exposure on the expression of genes regulating hormonal function, we analyzed a panel of candidate genes associated with steroidogenesis and hormonal signaling, including *Star*, *Cyp11a1*, *Cyp17a1*, *Cyp19a1*, *Hsd17b1*, *Hsd17b3*, *Hsd3b1*, and *Scarb1*.

In the F1 generation, thia exposure led to a significant upregulation of *Cyp11a1* (p = 0.02), *Cyp17a1* (p = 0.001), *Cyp19a1* (p = 0.02), *Hsd17b1* (p = 0.0004), and *Hsd3b1* (p = 0.003). In addition, *Star* expression showed a 1.94-fold increase, although this did not reach statistical significance.

In the F3 generation, we observed continued upregulation of *Cyp11a1* (p = 0.001), *Cyp19a1* (p = 0.02), *Hsd17b1* (p = 0.007), *Star* (p = 0.03), and *Scarb1* (p = 0.02), suggesting potential transgenerational effects on genes regulating steroid hormone biosynthesis.

### Immunofluorescence Analysis of CYP19A1 and ESR1 Proteins

To reveal the level of gene expression at the protein level, we performed immunofluorescence staining for CYP19A1 (aromatase) and ESR1 (estrogen receptor alpha) on ovarian tissue sections from F1 adult mice. Quantitative analysis of fluorescence intensity revealed a significant reduction in the expression of both proteins following gestational exposure to Thiacloprid, which is consistent with reduced levels of estradiol. Specifically, CYP19A1 staining was markedly decreased in granulosa cells of growing follicles (p=0.04), at the same time, ESR1 expression was also reduced in both granulosa and stromal compartments (p=0.01) (Fig.6). The observed discrepancy between mRNA and protein levels suggests possible post-transcriptional or translational regulation mechanisms disrupted by prenatal thiacloprid exposure, that led to reduced production of the estradiol hormone.

**Fig 6.**
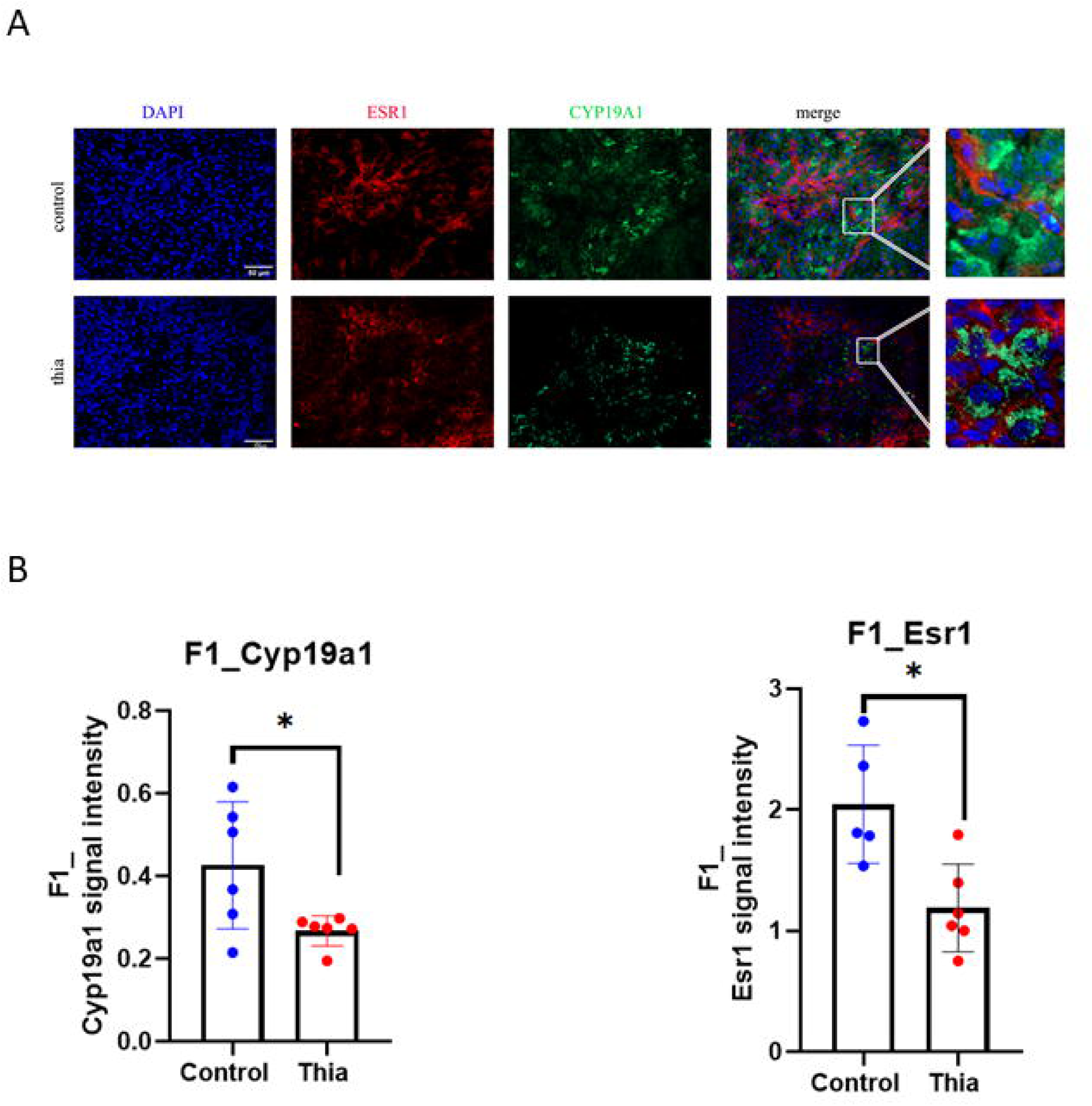
Immunostaining of CYP19A1 and ESR1 in Control and Thia-Exposed Ovaries. (A) Representative images of control and Thia-exposed ovaries. (B) Quantification of CYP19A1 and ESR1 in F1 generation. Statistical analysis shows p-values of 0.04 and 0.01, respectively.

These findings suggest that prenatal exposure to thia leads to long-lasting changes in the expression of genes critical to ovarian function. The alterations observed, particularly in genes involved in hormonal signaling, highlight the potential for thia to induce alterations in hormonal production and signaling. We suggest that epigenetic modifications at GRR genes may contribute to these alterations across generations.

### DNA methylation analysis in sperm of F1 and F2

In this study, paternal exposure impacts the female reproductive system development in F3. Sperm DNA methylation could also be involved in epigenetic control of gene expression. In our previous study, we analyzed the effects of thiacloprid on sperm DNA methylation in F1 to F2 generations [26]. Since females of F3 were obtained from the father of F2 generation, we looked closer at DNA methylation at genes studied in this study. We found that changes in 7 regions could have an impact on ovarian function (Fig. 7A-B). For example, we found a decrease in DNA methylation at the *Hormad1, Stk31, Tdrd1, Brdt, Ddx4 and Dazl and Inha* (Figure 7C, suggesting that increased gene expression in F3 females could be due to reduced DNA methylation in F1 farthers, which were transmitted to F2 and that could contribute to female of F3 generations.

**Fig 7.**
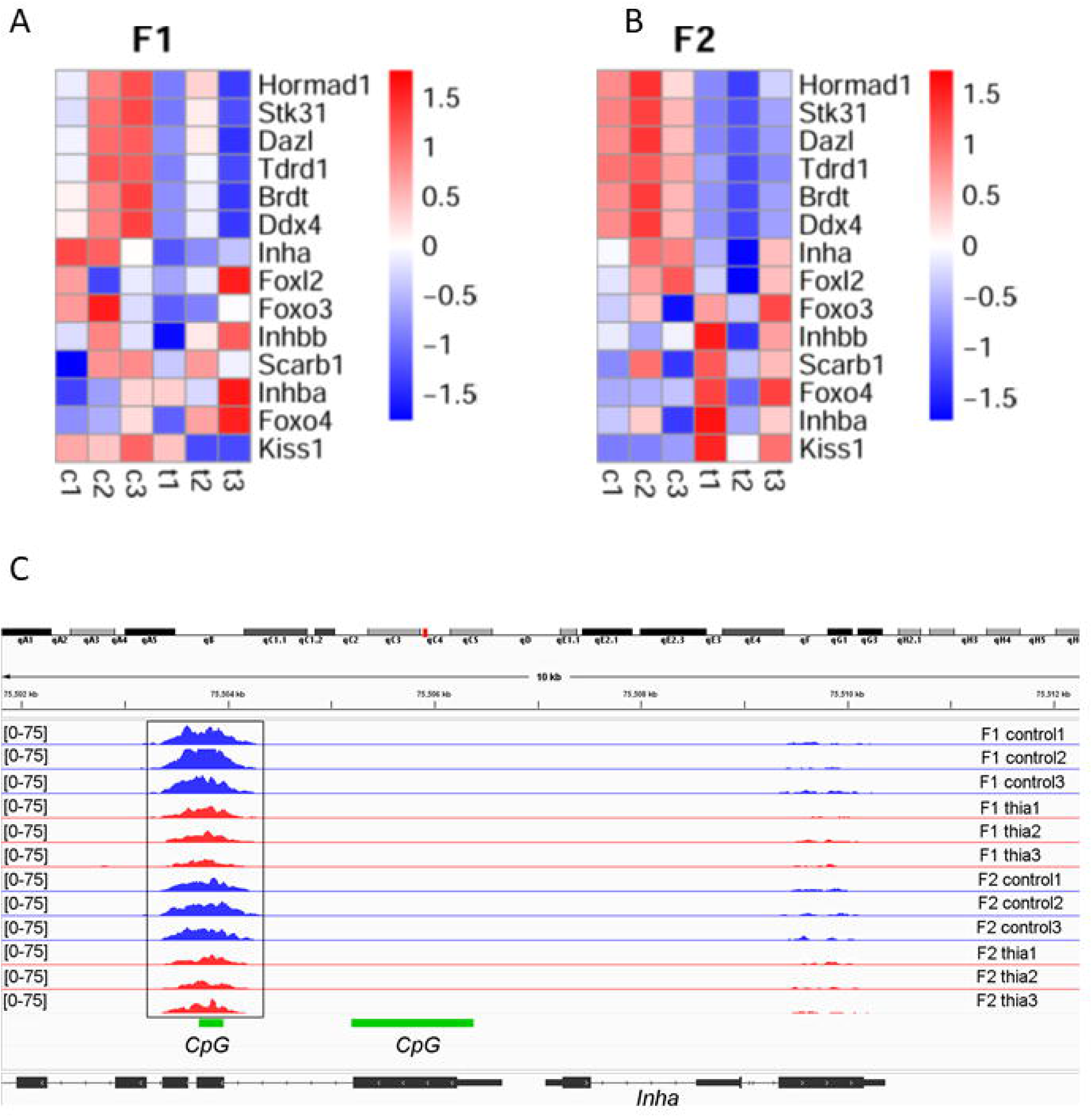
Heatmap of sperm DNA methylation in F1 and F2 generations. (A) Heatmap of F1 sperm DNA methylation (B) Heatmap of F2 sperm DNA methylation (C) Paternal Sperm DNA Methylation Profiles at the *Inha* Locus in F1 and F2 Generations Following Thiacloprid Exposure.

Thus, DNA methylation in paternal sperm was reduced in contrast to DNA methylation in female ovaries which was increased, suggesting that DNA methylation could be regulated in both sexes by different mechanisms.

## Discussion

### Impact of thia on ovarian weight and folliculogenesis

This study highlights the transgenerational effects of paternal exposure to thiacloprid (thia) on female reproductive health. We observed both metabolic and ovarian alterations in F1 and F3 generations, suggesting that paternal gestational exposure can have lasting impacts.

Although body weight remained unchanged in F1 females, a significant decrease was observed in F3 females (p = 0.018), indicating a delayed effect of exposure. This aligns with findings that prenatal exposure to environmental chemicals can cause transgenerational metabolic disturbances [6]. The ovary-to-body weight ratio showed opposite trends: increased in F1 (p = 0.01) and decreased in F3 (p = 0.05). The increase in F1 may reflect thyroid hormone disruption, as we previously showed that thia alters circulating TSH and T4 levels through reduced DIO1 expression [25]. Thyroid hormones regulate ovarian development, and their imbalance may affect folliculogenesis and steroidogenesis. The decrease in F3 suggests cumulative hormonal or metabolic disruption affecting ovarian tissue mass, consistent with other endocrine disruptor studies [7].

Histologically, we found a significant increase in multi-oocyte follicles (MOFs) in F1 females (p = 0.04), along with a tendency for more atretic follicles, suggesting impaired folliculogenesis. This phenotype resembles effects observed with other insecticides such as chlordecone [11]. MOFs were not assessed in F3 due to limited follicular changes in F1 and the complexity of full ovarian analysis.

Overall, our findings support the idea that paternal exposure to thiacloprid can affect female reproductive traits across generations, possibly through epigenetic mechanisms such as DNA methylation or histone modifications [8,]. Further research is needed to confirm the functional consequences and clarify the molecular pathways involved.

#### The impact on estradiol production

Our study assessed the impact of gestational thia exposure on estradiol serum levels in the F1 and F3 generations. We found a significant decrease in estradiol levels in both F1 (p = 0.015), and in F3 females (p = 0.0001). These results suggest that this exposure leads to hormonal disruption not only in the directly exposed first generation (F1) but also in the third generation (F3), pointing to potential transgenerational effects of ancestral paternal exposure on endocrine disruption.

Estradiol plays a crucial role in regulating reproductive health, and changes in its levels can result in reproductive dysfunction and altered fertility [9]. The significant decrease in estradiol levels observed in both F1 and F3 females is consistent with other studies showing that exposure to endocrine-disrupting chemicals can affect ovarian steroidogenesis and estrogen production [10]. The fact that estradiol levels were significantly altered in both generations highlights the persistence of these effects across generations, indicating the potential for cumulative or transgenerational endocrine disruption.

These findings emphasize the importance of understanding the long-term and transgenerational effects of environmental contaminants on endocrine function. Future studies should focus on the underlying molecular mechanisms, such as epigenetic modifications and changes in gene expression, that could explain the observed disruptions in estradiol production in both F1 and F3 generations.

#### Impact of gestational exposure to Thia on mitosis

We examined the impact of gestational thia exposure on ovarian function across multiple generations. Our results show that this exposure led to a significant decrease in estradiol (E2) levels and an increase in multi-oocyte follicles (MOF) in F1 females. These findings suggest that thia may disrupt normal ovarian development, impairing hormone production and folliculogenesis, which could affect reproductive outcomes. The increase in MOF, a condition that results from improper follicular development, is consistent with reports indicating that exposure to endocrine-disrupting chemicals (EDCs) can lead to abnormal follicle structures and impaired ovarian function [13,14].

To further investigate the mechanism underlying these alterations, we analyzed mitosis using phosphorylated histone H3 levels as a marker. In F1 ovaries, we observed a significant increase in histone H3 phosphorylation (p = 0.04), suggesting that gestational exposure to thia may lead to mitotic defects. These findings are in line with previous studies indicating that exposure to EDCs can disrupt the normal progression of the cell cycle, particularly mitosis, in ovarian cells. This disturbance could result in defective cell division, which may contribute to the abnormal ovarian structures and impaired folliculogenesis observed in the F1 generation [15,16]. However, in F3 ovaries, the trend was not statistically significant (p = 0.27), this suggests that the effects of thia exposure may be less pronounced via paternal exposure. It is possible that compensatory mechanisms in the F3 generation, such as genetic adaptation or other forms of physiological regulation, may counteract the initial disruptions caused by thia to male parents. Alternatively, the lack of a significant effect in the F3 ovaries could reflect a generational attenuation of the thia-induced disturbance, potentially indicating a recovery of ovarian function after multiple generations of exposure.

#### Impact of gestational exposure to thia in DNA methylation

Our investigation into the epigenetic consequences of prenatal thiacloprid exposure revealed notable alterations in DNA methylation patterns of key germline regulator genes (GRRs) and transposable elements, suggesting potential disruptions in germline development and genomic stability.

In embryonic day 15.5 (E15.5) ovaries from the F1 generation, we detected increased DNA methylation at the promoters of *Brdt*, *Dazl*, *Ddx4*, and *Hormad1*. These genes are pivotal for a germ cell development, and their hypermethylation may impact their expression, potentially leading to compromised germline population. This observation aligns with previous studies indicating that environmental exposures can induce epigenetic modifications in GRRs, thereby affecting reproductive outcomes [17].

In adult F1 ovaries, a similar trend of increased DNA methylation was observed at *Hormad1* (*p* = 0.05). In the F3 generation, elevated methylation levels were noted in *Brdt*, *Stk31*, and *Tdrd1*. These findings suggest that thiacloprid-induced epigenetic alterations may persist across generations, potentially through transgenerational inheritance mechanisms. Such transgenerational epigenetic effects have been documented in other studies examining neonicotinoid exposures [12].

Additionally, we assessed the methylation status of LINE-1 (L1) transposable elements, whose repression is crucial for maintaining genomic stability. We observed increased L1 methylation in both F1 and F3 generations (*p* = 0.05 for each), indicating potential interference with normal epigenetic reprogramming processes. Aberrant methylation of transposable elements like L1 has been implicated in genomic instability and has been proposed as a biomarker for environmental exposures [18].

Collectively, these findings suggest that prenatal exposure to thiacloprid may induce epigenetic modifications in GRRs and transposable elements, potentially disrupting germline development and compromising genomic integrity. The persistence of these epigenetic changes into subsequent generations underscores the need for further research to elucidate the mechanisms underlying these alterations and their implications for reproductive health.

#### Impact of thia on mRNA Expression

Our study provides evidence that prenatal exposure to thia significantly alters the expression of genes involved in gonadal and reproductive regulation in the ovaries of adult mice. Specifically, our RT-qPCR analysis demonstrated increased expression of key genes such as *Inhba* and *Fst* in the F1 generation, along with a more widespread upregulation in the F3 generation, including genes like *Inha*, *Inhbb*, *Foxl2*, and *Foxo3*. These findings suggest that prenatal exposure to thia can lead to lasting changes in ovarian gene expression, potentially disrupting normal ovarian function across generations.

The observed alterations in genes related to estrogen signaling are of particular concern, as these pathways play a crucial role in regulating ovarian development and function. The upregulation of *Inhba* and *Fst* in F1 suggests a potential disruption of follicular development. These genes are involved in follicle growth, and their altered expression could interfere with normal reproductive processes [19,20]. Notably, *Fst* plays a critical role in ovarian function by inhibiting activin signaling, thereby regulating follicle-stimulating hormone (FSH) secretion and folliculogenesis [21].

In the F3 generation, the upregulation of *Inha*, *Inhbb*, *Foxl2*, and *Foxo3* further supports the hypothesis that prenatal thia exposure results in long-term disruptions of ovarian function via paternal germline. *Foxl2* and *Foxo3* are transcription factors that are vital for the maintenance of ovarian reserve and the regulation of follicular development [22]. The observed changes in the expression of these genes, alongside the persistent effects in the F3 generation, suggest that thia-induced modifications may persist transgenerationally, possibly through epigenetic alterations of paternal germline.

The analysis of the steroidogenesis and hormonal signaling RT-qPCR of F1 reveals an increase of CYP11A1, CYP17A1, CYP19A, Hsd17b1, Hsd3b1, and a tendency to increase of Star, with a more widespread upregulation in the F3 generation. This upregulation suggests a disruption of the normal endocrine regulatory mechanisms, potentially leading to altered synthesis of sex steroids such as testosterone and estradiol. The increased expression of CYP11A1 and CYP17A1 indicates an enhanced conversion of cholesterol to pregnenolone and subsequent androgen precursors, while elevated CYP19A1 (aromatase) suggests a shift toward estrogen biosynthesis [28,29,30].

Such changes in steroidogenic gene expression may reflect a compensatory response to endocrine-disrupting stimuli during early development, or a direct effect of environmental contaminants on the hypothalamic–pituitary–gonadal (HPG) axis. Similar findings have been reported in animal models exposed to endocrine-disrupting chemicals, such as neonicotinoids or other xenobiotics, where increased transcription of steroidogenesis-related genes was associated with reproductive and neuroendocrine dysfunction [31,32].

Notably, these gene expression changes in the F1 generation may have long-term implications for reproductive capacity, sexual differentiation, and even behavior, especially if the exposure occurred during sensitive developmental windows [33].

The persistence of these alterations in gene expression across generations highlights the potential for epigenetic reprogramming induced by thia exposure. Epigenetic modifications, such as DNA methylation and histone modification, can lead to lasting changes in gene expression without altering the underlying DNA sequence [8]. Our findings align with previous studies suggesting that environmental toxins, including pesticides, can induce epigenetic changes that persist across generations and impact reproductive health [23,24].

In conclusion, our data provide compelling evidence that prenatal exposure to thiacloprid leads to significant and long-lasting changes in ovarian gene expression. These alterations, particularly in genes involved in estrogen signaling and oxidative stress response, may contribute to reproductive dysfunction in both the exposed generations and subsequent ones. Given the growing use of neonicotinoid pesticides, further research is needed to better understand the underlying mechanisms of these epigenetic modifications and their potential impact on reproductive health.

#### Decrease of CYP19A1 and estradiol level

Aromatase is responsible for the conversion of androgens into estrogens, and ESR1 mediates estrogen signaling in multiple tissues, including reproductive organs and the brain [34,35].

Our results revealed a marked decrease in fluorescence intensity for CYP19A1 and ESR1 in the F1, suggesting a reduction in protein expression or altered cellular localization. This observation contrasts with the increased mRNA levels of steroidogenic enzymes and may indicate post-transcriptional regulation, impaired protein stability, or feedback inhibition mechanisms. The concordant decrease in both aromatase and estrogen receptor proteins may contribute to the reduced estradiol levels detected in serum, and potentially to downstream physiological or behavioral effects.

These findings highlight a decoupling between transcriptomic and proteomic responses, pointing to a complex regulatory disturbance of the estrogen axis. This disruption may underlie early and potentially heritable impairments in endocrine homeostasis, warranting further investigation into its developmental and transgenerational consequences.

#### Paternal Epigenetic Transmission and Its Impact on Female Reproductive Development

In this study, we demonstrated that paternal exposure to thiacloprid can significantly impact the development of the female reproductive system, particularly in subsequent generations. The mechanisms underlying such effects are increasingly being attributed to epigenetic modifications in the paternal germline, notably changes in sperm DNA methylation patterns, which can persist across generations and influence gene expression in offspring [36,37].

In our previous work, we identified differentially methylated regions (DMRs) that persisted transgenerationally [26]. In the present study, we looked closer on DNA methylation in sperm across seven genomic regions, including *Hormad1, Stk31, Tdrd1, Brdt, Ddx4, Dazl, and Inha* (Figure 7A–C), all of which play crucial roles in germ cell development, meiotic progression, and ovarian physiology [38,39,40].

Interestingly, these genes exhibited hypomethylation in paternal sperm but hypermethylation in the ovaries of F3 females, suggesting a nonlinear and sex-specific epigenetic response. This observation implies that additional regulatory mechanisms may be active in the ovary, potentially as compensatory processes to counteract disrupted signals inherited via the male germline. Such mechanisms could include histone modifications, non-coding RNAs, or epigenetic crosstalk between DNA methylation and transcriptional repressors [41,42]. The reciprocal pattern of methylation between sperm and ovary points to a complex epigenetic reprogramming process occurring during early embryogenesis, which may not completely erase or override the environmentally induced marks.

Overall, these findings support the concept of paternal epigenetic inheritance and highlight the sensitivity of the female reproductive system to ancestral environmental exposures. They also raise concerns about the long-term consequences of pesticide exposure on reproductive health, even in the absence of direct contact with the chemical across generations.

## Methods

### Ethics Statement

All experimental procedures involving animals were authorized by the Ministry of National Education and Research of France (Authorization Number: APAFIS#17473-2018110914399411 v3). The animal facility used for this study is licensed by the French Ministry of Agriculture (Agreement Number: D35-238-19). All procedures adhered to the ethical principles outlined in the Ministry of Research Guide for the Care and Use of Laboratory Animals and complied with the ARRIVE guidelines.

Euthanasia procedures were conducted in accordance with Annex IV of Law 2013-118 issued by the Ministry of Agriculture, Food, and Forestry of France (February 1, 2013). Most animals were euthanized using a carbon dioxide (CO2) chamber. For hormone measurement analysis, mice were anesthetized with 130 mg/kg body weight of ketamine (Virbac, France) and 13 mg/kg body weight of xylazine/Rompun (Elanco, France) before blood collection via cardiac puncture, followed by euthanasia via decapitation.

### Animal Treatments and Dissections

Pregnant outbred Swiss female mice (RjOrl, Janvier Labs) were treated with thiacloprid (0 and 6 mg/kg/day) administered via oral gavage from embryonic day 6.5 (E6.5) to embryonic day 15.5 (E15.5). This treatment period corresponds to the germline lineage establishment window. The doses were determined based on the authorized daily dose for humans of imidacloprid (0.06 mg/kg/day) established by the French Agency for Food, Environmental and Occupational Health and Safety (ANSES), with doses set 100-fold higher to evaluate potential dose-dependent effects. Considering that mice tolerate higher doses, we recalculate the dose to human equivalent using a conversion index from [45]. Thus, the dose of 6ug/kg /day is ∼8 times higher than the human equivalent of the daily authorized dose, suggesting that the used dose is quite low.

Thiacloprid (Fluka, R1628-100MG) was suspended in olive oil and administered in a volume of 150 µL per mouse. Control mice received the same volume of olive oil. Treated and control pregnant females are referred to as the F0 generation. Their offspring (F1) were cross-bred with untreated, non-littermate mice to produce the F2 generation, and F2 males were crossed with untreated females to produce the F3 generation.

A minimum of 10 unrelated pregnant females were treated per dose, and 4–10 males from different litters were used for each assay. F1 and F3 males and females were euthanized at 60 days of age, corresponding to sexual maturity. At this stage, ovarian and spermatozoa samples were collected for analysis. This approach ensured sufficient sample sizes and biological replicates for statistical reliability.

### Morphology Analysis and Quantification of Follicles

Ovaries from F1 and F3 mice were dissected, fixed in Bouin’s solution, washed in PBS, and dehydrated before embedding in paraffin. Five-micrometer sections of the entire ovaries were cut, with every 5th section placed on a slide. The sections were deparaffinized using a Shandon Varistain 24–3 stainer, followed by immersion in distilled water for 1 minute.

Next, slides were stained with hematoxylin for 4 minutes and rinsed in running tap water for 1 minute. They were then washed four times in distilled water, followed by immersion in 100% alcohol containing 1% acetic acid for 15 seconds, and five washes in distilled water. The slides were treated with 3% ammonia (NH₄OH) for 1 minute, followed by five additional washes in distilled water. Afterward, slides were immersed in eosin for 2 minutes and dehydrated through a series of ethanol washes (two times in 70%, five times in 95%, and 1.5 minutes in 100%). Finally, slides were cleared with xylene (1 minute in xylene 1 and 2 minutes in xylene 2), dried for a few minutes, and mounted with Eukitt mounting medium.

The stained slides were scanned using the Hamamatsu NanoZoomer slide scanner, and follicles were quantified using the NDPview software. Follicles were counted when the oocyte nuclei were visible. The following classifications were used:

- **Primordial follicles**: oocytes surrounded by a single layer of flattened granulosa cells.
- **Primary follicles**: at least one granulosa cell of the single layer became cuboidal.
- **Secondary:** two layers of granulosa cells.
- **preantral follicles**: more than two layers of granulosa cells.
- **Antral follicles**: presence of an antral cavity.
- **Atretic follicles**: follicles containing at least two pyknotic granulosa cells.
- **Multi-Oocyte follicles**: follicles containing more than one oocyte.

On average, 74 sections per control sample and 75 sections per thiacloprid-treated sample were analyzed in the F1 generation.

### Estradiol Hormone Measurement by ELISA

Blood samples were allowed to clot at room temperature for 15–30 minutes. After clotting, the samples were centrifuged for 10 minutes in a refrigerated centrifuge to separate the serum. The resulting supernatant was aliquoted and stored at −80°C until further use.

Estradiol levels were measured using the Mouse Estrogen ELISA Kit (96-Test, orb408998-96, CliniSciences), following the manufacturer’s instructions. Data were averaged, and estradiol concentrations were presented as pg/ml.

### Immunofluorescence on paraffin sections

Epitopes were unmasked by heating the tissue sections in 0.01 M citrate buffer (pH 6) at 80°C for 20 minutes. After blocking with 4% BSA in 1X PBS containing 0.05% Tween (PBS-T), the sections were incubated overnight at 4°C in a humidified chamber with the following primary antibodies: mouse anti-MSY2 (1:200) (Sc-393840, Santa Cruz) and rabbit anti-phosphohistone H3 ser10 (1:200) (07-327, Millipore).

After washing in PBS containing 0.04% Kodak Photo Flo solution, sections were incubated with the appropriate Alexa Fluor-conjugated secondary antibody (1:500, Invitrogen) for 1 hour at room temperature in a humidified chamber. The sections were then mounted using Vectashield mounting medium containing 0.001% (v/v) 4,6-diamidino-2-phenylindole dihydrochloride (DAPI) for nuclear staining.

Images were captured using an AxioImager microscope (Zeiss, Le Pecq, France) equipped with an AxioCam MRc5 camera and AxioVision software version 4.8.2, using a 20X or 40X objective lens

### DNA extraction

DNA extraction was performed using a DNeasy Blood & Tissue kit (69506; QIAGEN). First, embryonic and adults’ ovaries were physically disrupted with a tissue lyser (QIAGEN in AL buffer) using Tungsten Carbide Beads (69997; QIAGEN). DTT (10 mM) was added to the lysis solution. Incubation with lysis solution was performed at 56°C overnight. The DNA extraction protocol included an RNase A (19101; QIAGEN) treatment step to eliminate contaminating RNA. The DNA concentration was measured using the QuantiFluor dsDNA system (E2670; Promega). The quality of the DNA was assessed by running samples on a 0.7% agarose gel; a homogenous high-molecular weight signal was observed for each sample. DNA extraction from embryonic ovaries was performed according to DNeasy Blood & Tissue kits (69506; QIAGEN) without modifications.

### Methylated DNA precipitation (MEDIP) and MEDIP–qPCR

For DNA methylation analysis in embryonic and adult ovaries, the EpiMark Methylated DNA Enrichment Kit (#E2600S; NEB) was used. A total of 6,000 ng of DNA or 3,000 ng (embryonic ovaries) was sonicated using a Qsonica sonicator with the following parameters: efficiency, 60%; total sonication time, 8 min, 20 s “on” and 20 s “off.” The average size of sonicated DNA was 300 bp. The sonicated methylated DNA was precipitated using MBD2a-protein A-coated beads, the methylated DNA-MBD2a-protein A–coated bead complex was washed, and the DNA was eluted with elution buffer. The DNA concentration was determined according to the level of fluorescence produced by a dsDNA-binding dye (Promega). ∼3–4 ng (embryonic ovaries) of methylated DNA was recovered after precipitation, and the unprecipitated sonicated starting material was used as the input.

For MEDIP–qPCR, enriched DNA and input were diluted to equal concentrations, and qPCR was performed. Primers were designed using the coordinates of differential peaks (Table S1). The sequences were retrieved from USCSC repeat masks, and Primer-Blast was used to design primers. Equal amounts of enriched DNA and input were used for PCR. The background was normalized to the *Rplp0* unmethylated region. We used seven replicates for the control and seven for *thia for adults’ ovaries and n=5 for control and thia for E 15.5 ovaries*, and each replicate contained DNA from E15.5 ovaries. The data for each gene were normalized and compared with the control.

### RNA extraction and RT-qPCR

Ovaries were stored at −80°C until use. RNA extraction was performed using the **RNeasy Plus Mini Kit** (Qiagen, 2,552,951) on 60-day-old mice from the control (“0” group) and the “6 mg/kg/day” treatment group. Approximately 10 mg of ovarian tissue was used per sample.

Ovary samples were lysed and homogenized using a TissueLyser (Qiagen) with 5 mm stainless-steel beads (Qiagen, 69,989). DNA was removed by passing the lysate through a DNA elimination column. To provide optimal binding conditions, one volume of 70% ethanol was added to the lysate. The RNA was then bound to the RNeasy silica membrane, followed by an additional treatment with DNase using the RNAse-free DNase set (Qiagen, 79,254) directly on the column.

RNA was washed with RW1 and RPE buffers from the Qiagen kit to remove impurities and eluted in 50 µL of RNase-free water. For RT-qPCR, eight biological replicates from the control group and eight from the treated group were used for each group.

Reverse transcription (RT) was performed using 1 µg of total RNA and the iScript™ cDNA Synthesis Kit (BioRad, 1,708,891), following the MIQE guidelines. *Rpl37a* was used as housekeeping genes, as they showed no variation across replicates in our RNA-seq data. Data were presented as fold changes relative to control ± SD.

Primers for this study were selected using the Primer-BLAST tool (NCBI, ncbi.nih.gov) and were designed to span exon-to-exon junctions. The primer sequences are listed in **Supplementary Table**. Statistical significance was assessed using a nonparametric Mann– Whitney test.

### Statistical analyses

We used the minimum number of animals according to the requirements of the EU ethics committee. The number of animals used was specified for each experimental procedure. We performed non-parametric Mann–Whitney tests to assess statistical significance in qPCR experiments, immunofluorescence. Non-parametric Kruskal–Wallis test was used to assess the statistical significance in body weight measurements.

## Data availability

All data generated or analyzed during this study are included in this published article and its supplementary information files. Additional datasets or materials related to the study can be made available upon reasonable request.

## Acknowledgements

We would like to thank the H2P2 platform (UMS Biosit Inserm UMS 018, CNRS UMS3480) for assistance with the preparation and analysis of paraffin thyroid sections.

## Author information

These authors contributed equally: Ouzna Dali and Chaima Diba Lahmidi.

## Contributions

O.D., C.D.L., C.K., P.Y.K.: Methodology. F.S.: Investigation, editing, conceptualization, supervision, methodology, investigation, O.D and FS writing and draft preparation.

## Corresponding author

Correspondence to FS

## Abreviations

Brdt: Bromodomain testis associated
Cyp11a1: Cytochrome P450 family 11 subfamily A member 1
Cyp17a1: Cytochrome P450 family 17 subfamily A member 1
Cyp19a1: Cytochrome P450 family 19 subfamily A member 1 (aromatase)
*Dazl*: Deleted in azoospermia-like
Ddx4: DEAD-box helicase 4
DNA: Deoxyribonucleic Acid
E2: Estradiol
E6.5-15.5: embryonic day 6.5-15.5
EDCs: Endocrine-Disrupting Chemicals
Esr1: Estrogen receptor 1
F0: Founding Generation or Parental Generation
F1: First Filial Generation
F2: Second Filial Generation
F3: Third Filial Generation
*Foxl2*: Forkhead Box L2
*Foxo3*: Forkhead Box O3
*Foxo4*: Forkhead Box O4
FSH: follicle-stimulating hormone:
*Fst*: Follistatin
GRRs: Germ line reprogramming responsive genes
H&E: hematoxylin and eosin
Hormad1: HORMA domain containing 1
Hsd17b1: Hydroxysteroid 17-beta dehydrogenase 1
Hsd17b3: Hydroxysteroid 17-beta dehydrogenase 3
Hsd3b1: Hydroxy-delta-5-steroid dehydrogenase, 3 beta- and steroid delta-isomerase 1
*Inha*: Inhibin Alpha
*Inhba*: Inhibin Beta A
*Inhbb*: Inhibin Beta B
LINE-1: Long Interspersed Nuclear Element-1
Medip: Methylated DNA immunoprecipitation
MOF: multi-oocyte follicles
MSY2: Mouse Spermatogenic Factor 2
Ncor1: Nuclear receptor co-repressor 1
Rplp0: Ribosomal Protein Lateral Stalk Subunit P0
RT-qPCR: reverse Transcription quantitative Polymerase Chain Reaction
*Scarb1*: Scavenger Receptor Class B Type 1
*Star*: Steroidogenic Acute Regulatory Protein
Stk31: Serine/Threonine Kinase 31
Tdrd1: Tudor domain containing 1
Thia: Thiacloprid

